# Phenotypic plasticity promotes recombination and gene clustering in periodic environments

**DOI:** 10.1101/092700

## Abstract

The impact of changing environments on the evolution of genetic recombination is still unclear. While the Red Queen hypothesis provides a reasonable explanation for recombination, it requires coevolution with antagonistic species, such as host-parasite systems. We present a novel scenario for the evolution of recombination in changing environments: the genomic storage effect due to phenotypic plasticity. Using an analytic approximation and Monte Carlo simulations, we demonstrate that recombination evolves between a target locus that determines fitness, and a modifier locus that modulates the effects of alleles at the target. Evolution of recombination by this plasticity effect does not require antagonistic inter-specific interactions and, unlike in previous models, it occurs when only one target locus codes for a trait under selection. Furthermore, if the effects of multiple target loci are modified by the same plasticity locus, then the recombination rate among the target loci will tend to decrease, clustering the loci that influence a trait. These results provide a novel scenario for the evolution of recombination, highlighting the importance of phenotypic plasticity for recombination modification.

## INTRODUCTION

What governs the evolution of genetic recombination is a subject longstanding interest, with tremendous development and progress as well as outstanding questions. A genetic basis for recombination, such as response to selection (Korol and Iliadi 1994), genetic variation underlying recombination rates (Chinnici 1981, Charlesworth and Charlesworth 1985a and 1985b, Brooks and Marks 1986, Brooks 1988, Williams et al 1995, Kong et al 2002), and modifiers of rates of recombination (Ji et al 1999, Kong et al 2008), have been reported. However, our understanding of how recombination evolves comes primarily from theory. Theory suggests that recombination evolves in populations with negative linkage disequilibrium (negative LD, where alleles of opposite fitness effects at two different loci co-segregate). In such populations, recombination couples beneficial alleles at the two recombining loci and generates the fittest haplotype. Then, the recombination modifier allele, associated with the fittest haplotype, hitchhikes to a higher frequency, increasing the recombination rate in the population. In equilibrium populations, which are already at their optimum haplotypes, there is no advantage to reshuffling of the gene combinations and recombination is expected only to decrease, a phenomenon known as the reduction principle (Feldman et al. 1980, 1997).

In non-equilibrium populations, undesirable association between loci (negative LD) arises from a steady influx of mutations or from constantly changing environments. A steady influx of mutations repeatedly reintroduces diversity that would otherwise be depleted by selection. Then, under the directional selection, negative LD is generated either by negative epistasis or by chance (genetic drift). For epistasis to promote recombination, however, the interaction between the selected loci needs not only to be negative but also weak (Barton 1995). This narrows the range of parameters where recombination evolves under the negative epistasis model.

The conditions that permit the evolution of recombination under directional selection are greatly expanded in finite populations. Here, the interplay between drift and selection generates negative LD between two selected loci due to Hill-Robertson interference (1966). Hill and Roberson showed that the negative association between the selected loci impedes selection. Then, as the beneficial gene combinations (adaptive alleles at both of the loci) quickly fix, and detrimental combinations quickly perish, populations are left with the prevalence of mismatched pairs, i.e. negative LD. That recombination modulates effects of selection at the linked sites is well appreciated (e.g. McGaugh et al 2012), particularly given numerous reports of selective sweeps (Maynard Smith and Haigh 1974, Kim and Stephan 2002, Nielsen 2005) – if pervasive selection affects linked neutral variants so it does linked selected alleles. While the preponderance of negative LD under interference provides a plausible model for evolution of recombination irrespective of the form or presence of epistasis, population size, or even type of selection (for example selection against deleterious alleles, Keightley and Otto 2006), it is worth noting that in the absence of balanced polymorphism evolution of recombination still depends on steady influx of mutation.

Balancing selection in changing environments, on the other hand, might generate considerable diversity and constant fluctuating LD independent of influx of new mutants. Charlesworth (1976) was the first to describe the evolution of recombination when fluctuating epistasis arises from environmental changes, random or periodic. In particular, length of the periodic succession of environments (period) is a strong predictor of the optimal recombination rate (relatively fittest) under these scenarios. The optimal rate increases with shorter periods, provided the period exceeds two generations, and with the strength of linkage of the rate modifier to the selected pair of loci. Furthermore, Sassaki and Iwasa (1987) showed that the strength of selection seem to have very small effect on the optimal recombination rate under periodic environments, with very wide range of selection strength resulting in a rather narrow range of optimal recombination rates. While, at the time it appeared that this fluctuating epistasis mechanism (following Peters and Lively, 1999) might not be widespread, it was soon suggested that coevolution between antagonistic species, such as parasite and host, provides a simple mechanism for such changing environments (Hamilton 1980), a scenario referred to as the Red Queen hypothesis for the evolution of sex (Bell 1982).

The Red Queen hypothesis has become nearly synonymous with the evolution of recombination under changing environments (here, environments are the frequencies of pathogen and host genotypes). The Red Queen offers simultaneously two convincing conditions for evolution of recombination: 1) diversity at both of two selected loci due to negative frequency dependence arising from the coevolution between the antagonistic species, and 2) cycling linkage disequilibrium (LD) as novel rare haplotypes become advantageous and common ones become detrimental. Indeed, recombination seems to be more prevalent in presence of parasites in natural populations (review by Neiman and Koskella 2009). However, under the Red Queen hypothesis selection must be strong in at least one species (Salathé et al 2009) in order to assure appropriate cycling in fitness. Also, the Red Queen models assume that the two loci contribute to the interaction between competing species. In nature, however, parasites may evolve mechanisms to express a single antigen (i.e. allele) at the time (Donelson 1995, Barbour and Restrepo 2000, Kusch and Schmidt 2001). Finally, the Red Queen hypothesis requires interaction with an antagonistic species.

In this paper we study an alternative mechanism that can produce cycling LD between two loci: namely, LD between a plasticity (or robustness) modifier locus and its target locus, which can occur in the absence of antagonistic species, or constant influx of mutation. Such LD can occur when environments change periodically, due to a mechanism called the “genomic storage effect” (Figure 1, Gulisija et al 2016). The basic idea behind the genomic storage is that alleles can survive periods of adversity by escaping to a genetic background that ameliorates the effects of selection and stores diversity until conditions change. This novel model of balancing selection in periodic environments assumes sign epistasis: the plasticity allele is beneficial when paired with the detrimental allele (benefit of plasticity), but harmful when paired with an adaptive allele (cost of plasticity). Selection promotes association between the detrimental allele and the plasticity modifier and so it generates LD between the two loci; more over the magnitude of this LD is amplified by balancing selection (Figure S1). When the environment changes and the target allele becomes beneficial, the newly adaptive allele is still linked with now the costly plasticity allele, i.e. negative LD arises. This dynamics of changing LD, such that the best combinations of alleles are underrepresented when the target allele becomes beneficial could promote recombination.

**Figure 1.**
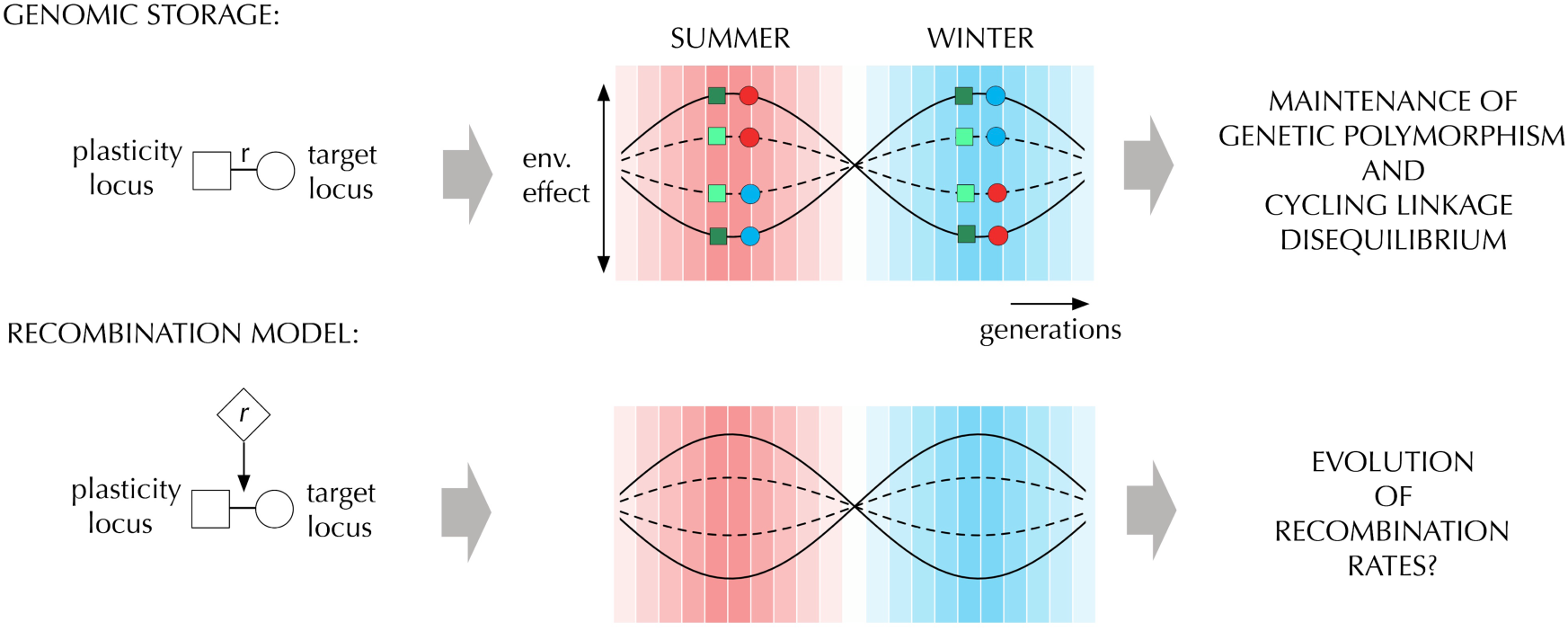
Schematic representation of the genomic storage effect and the recombination model. As environmental effects oscillate, the summer (red) and the winter (blue) alleles at the target locus recombine to a less harmful genetic background at the plasticity modifier locus. The plasticity modifier locus modulates the effect of selection at the target locus by making alleles at the target locus more (non-plastic, dark green) or less (plastic, light green) visible to selection. The dynamics between the two loci allows unfavorable alleles to be stored until conditions change, generating the genomic storage effect (Gulisija et al. 2016). The genomic storage effect generates both balanced polymorphism and cycling linkage disequilibrium. Here we study whether or not this type of cycling linkage disequilibrium can lead to the evolution of recombination.

The genomic storage effect can arise in populations that experience periodic environmental changes, such as annual seasonality. It is widely appreciated that phenotypic plasticity may mitigate adverse effects of selection in adverse or perturbed habitats (West-Eberhard 2003, Price 2006, Lande 2009). While plasticity might be result of an environmental sensitivity of the coding locus, empirical studies also suggest that plasticity might be modulated by sites other than the affected locus, also known as the epistatic model of plasticity (for a review see Scheiner 1993). Furthermore, mapping studies confirm many quantitative trait loci that modulate phenotypic plasticity in several model organisms (Stratton 1998; Leips and Mackay 2000; Bergland et al. 2008; Tetard-Jones et al. 2011). It is this epistatic model of plasticity that can give rise to the genomic storage effect described above. The genomic storage effect, however, is limited to either relatively large populations, or strong selection, or recurrent mutation. These conditions are likely to hold for populations subject to strong seasonality (see arguments given in Gulisija et al 2016). There, cycling linkage disequilibrium might induce selection for recombination between the plasticity and the target locus, and it might even bring to proximity multiple target loci whose effects are modulated by the same plasticity locus (i.e. co-modulated due to a shared pleiotropic transcription factor) as they would gravitate to the same recombination distance with the modifier.

In this study, we explore the evolution or recombination rates between a plasticity modifier and its target locus, as well as between multiple target loci that are controlled by the same plasticity modifier, under the conditions of environmental variability that generate the genomic storage effect (Gulisija et al. 2016). First, we conduct a stability analysis in the infinite-population limit to understand the deterministic dynamic at the recombination modifier locus. Then, we study the evolution of the recombination, plasticity, and target locus/loci in finite populations, via Monte Carlo simulations. We demonstrate that genomic storage leads to the evolution of recombination between the plasticity modifier locus and its target locus. Furthermore, when we study multiple target loci, we show that same effect leads to clustering of the target loci whose effects are modulated by the same plasticity locus, and therefore increases the magnitude of frequency oscillations and maintenance of diversity at the target loci.

## MODEL AND METHODS

To model the evolution of the recombination rate between a plasticity and target locus we expand the Wright-Fisher population model described in Gulisija et al (2016) to consider three loci: a recombination modifier, a bi-allelic plasticity modifier, and bi-allelic target locus whose fitness effects depend upon a periodically changing environment. At the target locus, an ancestral allele (*a*) is favored over the derived allele *(d)* for half of the environmental period, whereas *d* is favored over *a* for the other half of the environmental period. At the plasticity modifier locus (a epigenetic modifier or a transcription factor, for example), the plasticity modifier allele (*M*) alters the target locus phenotype so as to increase its fitness in detrimental environments (benefit of plasticity; West-Eberhard 2003, Lande 2009), but also decreases fitness in beneficial environments (cost of plasticity; DeWitt et al. 1998, Lande 2009). Then, the magnitude of fitness oscillations at the target locus are smaller in genotypes that carry *M* at the plasticity locus (as Gulisija et al. noted: also equivalent to action of a robustness modifier, de Visser 2003) as compared to the carriers of the non-modifier allele (*m*). The plasticity modifier locus and its target locus recombine at a rate controlled by a recombination modifier locus. The frequencies of resulting haplotypes are subject to deterministic effects of haploid selection in a constant population (soft selection, Wallace 1975) and of recombination between the three loci, and to the stochastic effects of genetic drift in finite populations, in each generation.

In this section, we first describe the deterministic dynamics: selection and recombination, when the two competing alleles at the recombination modifier locus are present. We undertake a stability analysis in the infinite-population limit, based on these deterministic dynamics. Finally, we describe the results of Monte Carlo simulations that allows for multiple recombination alleles, multiple target loci, and include the effect of genetic drift in finite populations.

### Selection

Combinations of alleles at the three loci (the recombination, plasticity, and target locus, with two competing alleles at each) form eight distinct haplotypes, *r*_1_*ma*, *r*_1_*Ma*, *r*_1_*md*, *r*_1_*Md*, *r*_2_*ma*, *r*_2_*Ma*, *r*_2_*md*, and *r*_2_*Md*, where *r*_1_ and *r*_2_ are recombination modifier alleles that produce different recombination rates between the plasticity and the target locus. The frequencies of the eight haplotypes in each generation are first modified by selection such that post-selection frequency of the haplotype is the product of its pre-selection frequency and its fitness:

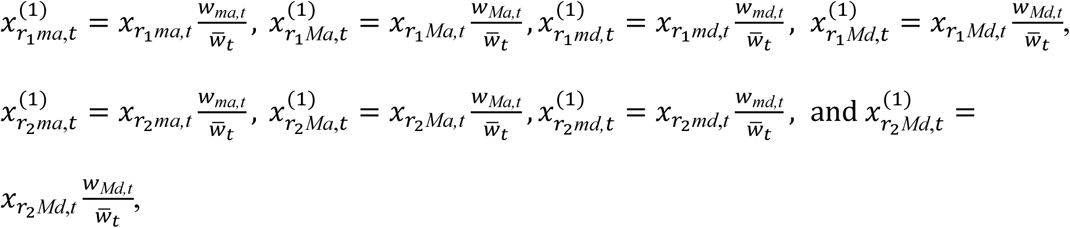

where

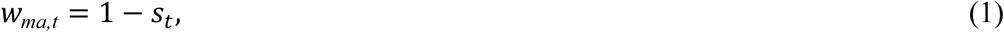

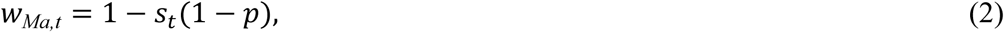

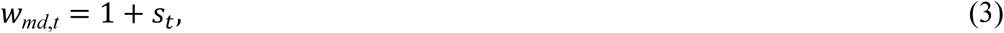

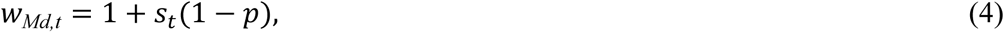

and

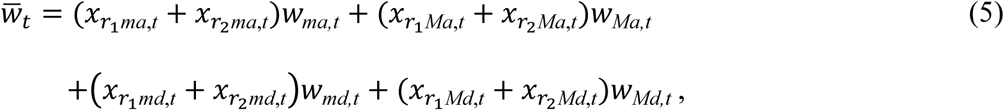

with

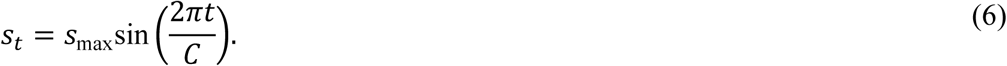

Here *s_t_* denotes the periodic environmental fitness effect at the target locus at the time *t*, which follows a sinusoidal function with maximum at *s*_max_ over a period of *C* discrete generations.

### Recombination

The post-selection haplotype frequencies are subsequently modified by recombination between the three loci, assuming an additive recombination phenotype between the two competing alleles at the recombination locus. Therefore, the two chromosomes recombine with the rate *r*_1_ or *r*_2_ if the carry the same allele, and with the rate *r_c_*= (*r*_1_ + *r*_2_)/2 if they carry different alleles. (Note that this does not mean additive in fitness, as an intermediate phenotype might carry an advantage or disadvantage compared to the both of the recombination phenotypes.) The physical map of the three loci is assumed (without loss of generality) to be recombination – plasticity – target, and the recombination modifier and the plasticity locus recombine at a fixed rate, *R*. Then, the haplotype frequencies following recombination are

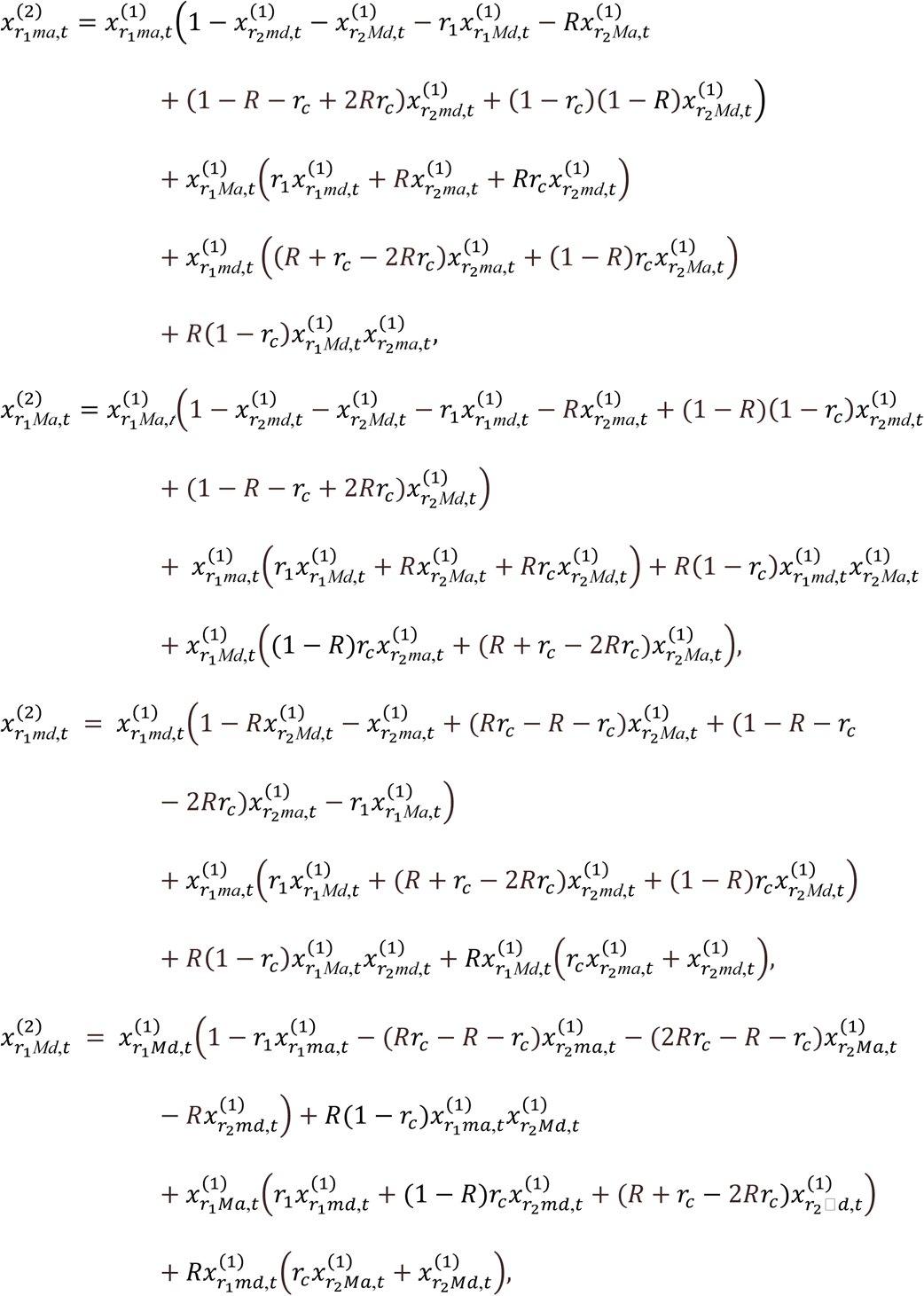

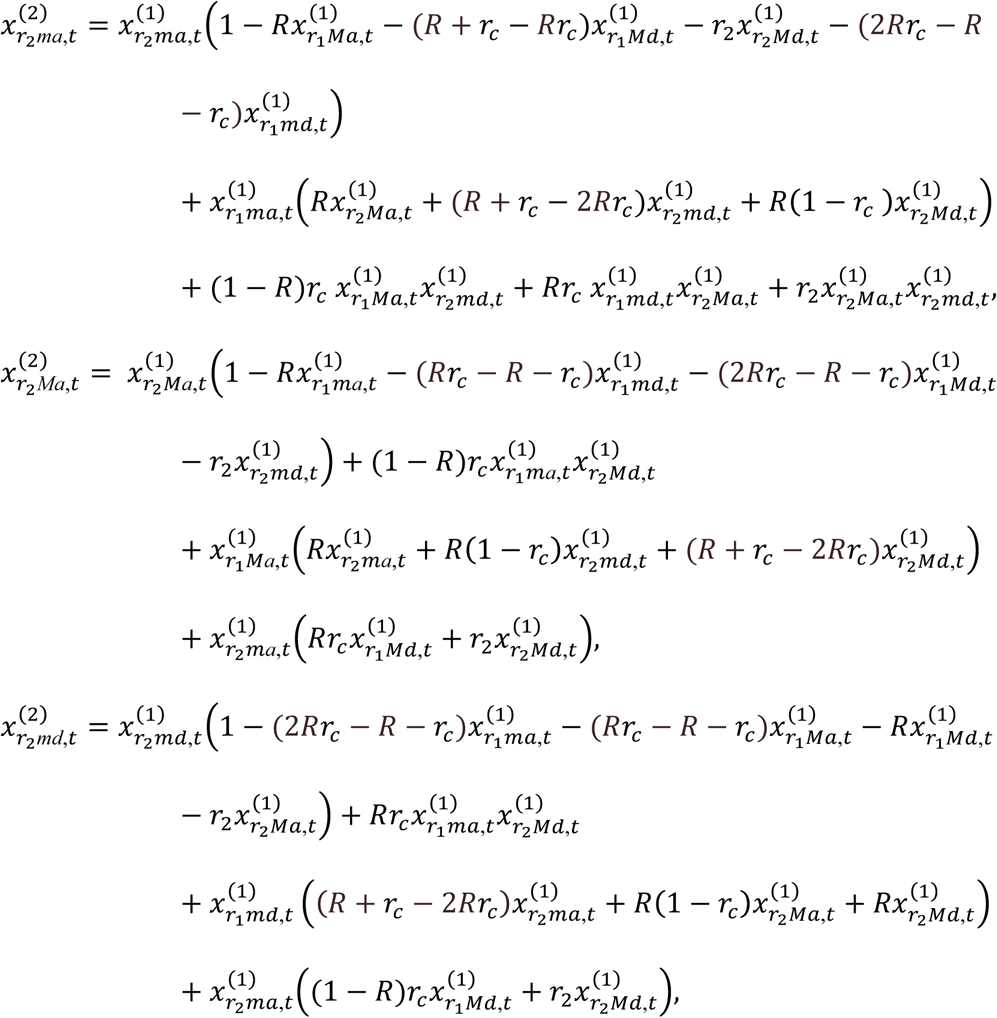

and

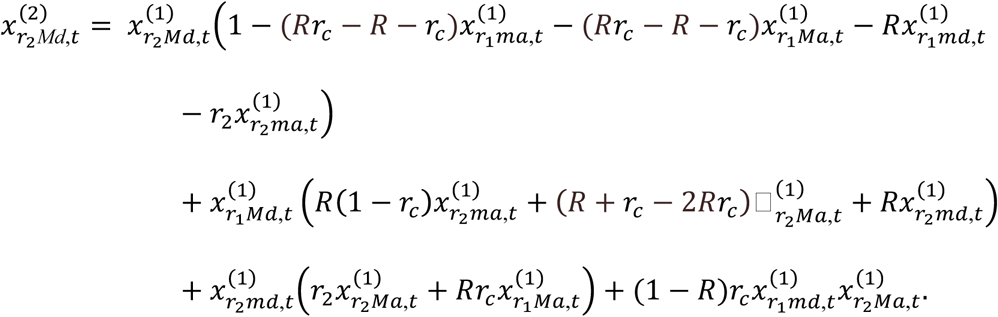

In the absence of genetic drift, these frequencies become the starting allele frequencies in the subsequent generation; whereas in the presence of drift the subsequent generation is formed by sampling with replacement from these frequencies.

### Stability Analysis

We first analyze this model in the infinite-population limit, neglecting both genetic drift and mutation. To do so we numerically evolve the discrete-time frequency equations to identify each equilibrium that is polymorphic at both the plasticity and target locus (defined here as the case when minor allele frequency does not fall bellow 0.01), irrespective of the frequency of alleles at the recombination locus. We focus on such polymorphic equilibria because, in the absence of influx of mutation, recombination evolves only if balanced polymorphism is present at the two loci (i.e. LD is maintained by selection). Next, for each such equilibrium identified numerically we compute the Jacobean matrix of the deterministic system over a full period of fitness oscillations and the corresponding leading eigenvalue to determine if the equilibrium is locally stable or not.

We studied the deterministic dynamics across the all possible pairwise combinations of recombination rate alleles (to two-digit precision), with environmental periods *C* = 10, 20, 40, or 80 generations. As Sassaki and Iwasa (1987) showed, optimum recombination rate is rather robust to the selection strength in the periodic environments, hence we do not vary the maximum environmental effect size, keeping *s*_max_ = 0.1. Although genomic storage can occur across various strengths of the plasticity effect (Gulisija et al 2016) for simplicity we fix the parameter *p* = 1. However, later we relax this assumption for finite-population simulations. Finally, Charlesworth (1976) noted that there is no optimum recombination rate for a given periodic environment, but the optimum rate increases with stronger linkage of the recombination modifier to the selected loci (*R*). Therefore, we set *R* = 0.5 (unlinked recombination modifier) as this will produce a lower bound on the range of recombination rates that might evolve under the genomic storage, making our conclusions about the evolution of recombination conservative. We initiate allele frequencies at each of the three loci ranging from 0.05 to 0.95 (in 0.1 increments), and we evolve the deterministic system for at least 1000 generation, and until either the plasticity or target locus fixes (frequency drops below 10^−4^) or until the same sequence (up to 8 digit accuracy) of haplotype frequencies is repeated in two consecutive environmental cycles, which we consider an equilibrium outcome.

### Monte Carlo Simulations in Finite Populations

To confirm that recombination modifier can invade in a non-recombining finite population, i.e. in the presence of genetic drift, and to allow recurrent mutations across the range of recombination phenotypes (*r* ~ U[0, 0.5]), we conducted Monte-Carlo simulations. We start each simulation in a monomorphic population at the three loci: with no recombination between the plasticity and the target locus, with allele *m* at the plasticity modifier, and with allele *a* at the target locus. Mutation randomly introduces diversity at each of the loci with chance *Nμ* = 0.1 per generation. At the recombination modifier locus, the mutant recombination rate is randomly chosen from a uniform distribution, U[0, 0.5]. The other two loci reversibly mutate between alleles *m* and *M*, or alleles *a* and *d*. Mutation is followed by recombination and sampling with replacement (drift). Each of the two parents is randomly sampled (with replacement) and retained for reproduction proportional to its fitness relative to the maximum fitness in the population. The plasticity and the target locus recombine between the two parental chromosomes, with probability depending on the alleles they carry at the recombination modifier in an additive manner as described above. The recombination modifier locus in the two gametes recombine with the plasticity – target sequence with probability *R*. Each pair of parents is chosen from the population sequentially until the next generation of *N* individuals is assembled. Simulations run for a burn-in duration of 100*N* generations (until genetic variance at the target and the plasticity locus and the stationary distribution at the recombination locus stabilize) and for additional 100*N* generations during which we record the stationary distribution of recombination rates (to two decimal place precision).

Gulisija et al (2016) showed that the genomic storage effect promoting balanced polymorphism at the target and plasticity loci increases with population size or *s*_max_, and with mutation rate. We studied populations of size *N* = 25,000 with a relative large value of *s*_max_, corresponding to strong selection from environmental variation. We obtained the stationary distribution of recombination rates between the plasticity and the target locus, *r*, under the cycle of fitness oscillations *C* = 10 with *s*_max_ = 0.25 and 0.5, *C* = 20 with *s*_max_ = 0.125 and 0.25, *C* = 40 and 80 with *s*_max_ = 0.15 and 0.075, all under *p* = 1, in an ensemble of over one thousand Monte Carlo simulations. To demonstrate that recombination also evolves with weaker absolute selection in larger populations, we also conducted simulations based on the deterministic recursion given earlier (competing two recombination rates), but with the multinomial sampling (reproduction/drift) of haplotype frequencies. Here, we examined the effects of *N* = 10^5^ or 10^6^, and *p* = 0.5 with *s*_max_ = 0.01, 0.02, or 0.03, in over forty-thousand replicate simulations.

#### Two target loci

We also study a genetic system similar to the one above, but with two target loci whose alleles each contribute additively to fitness, and whose effects are modulated by a single plasticity modifier locus. In this context we are especially interested in the evolution of the recombination rate among the target loci themselves. We study two particular scenarios: simultaneous evolution of the recombination rates between the plasticity and the first target locus and between the two target loci, and a sequential scenario. In the first scenario, a population migrates to the new periodic habitat and the two recombination rates are each initially monomorphic and drawn from U[0, 0.5], and then subsequently evolve. In the second scenario, a population starts in equllibirum based on the three-locus model above, and a new target locus in introduced, as for example may occur due to gene duplication and subfunctionalization (Stoltzfus 1999, Force et al 1999). In the second scenario the initial target locus recombines with the plasticity modifier with optimal recombination rate, whereas the newly arisen target locus has a random initial recombination rates with the existing target locus, drawn from U[0,0.5]), and it is placed either between the plasticity modifier and the first target locus (for *C* = 10 or 20) or further downstream of the first target locus (*C* = 10, 20, 40, or 80). In both scenarios we assume additive contribution to fitness of the multiple target loci. We postulate that if the two target loci are controlled by the same plasticity locus, then the target loci will evolve to the same recombination rate with the joint plasticity locus and thus cluster together. Additionally, in the simultaneous rate co-evolution model we also examine the distance between target loci even when they would not gravitate to the same recombination to the plasticity locus, i.e. not equally distant recombination modifiers.

We obtain the distribution of recombination rates between the two target loci, *r*’, under the cycle of fitness oscillations *C* = 10 with *s*_max_ = 0.25 and 0.5, *C* = 20 with *s*_max_ = 0.125 and 0.25, *C* = 40 and 80 with *s*_max_ = 0.15 and 0.075, all under *p* = 1. The simulations times are as above.

#### Multiple target loci

A model with more than two target loci may exhibit qualitatively different behavior that the case of only two target loci. For a small set of parameters (with *C* = 20), we explore the evolution of polymorphic clusters of loci acting in unisom (i.e. supergenes, e.g. Joron et al 2011, Wang et al. 2013). Again we consider two scenarios: we conduct a three target loci simulation where three recombination evolve simultaneously, and *n* target loci model where the recombination rate evolves between a cluster of *n*-1 target loci at equilibrium under the genomic storage and a *n*^th^ locus introduced sequentially (*n* = 3, 4, 5, 6, 7, or 8; *n* > 3 with *s*_max_ = 0.075). These simulations are conducted for 100*N* burn-in generations until stationary distribution is reached, and another 10,000 generations during which we record the stationary distribution of the recombination rate among target loci.

## RESULTS

The genomic storage effect leads to evolution of recombination between the plasticity modifier locus (such as an epigenetic modifier or a transcription factor) and the target locus or loci whose fitness effects are modulated by the modifier. At the same time, genomic storage favors complete linkage between two target loci controlled by the same modifier locus, that is co-modulated target loci. Within these clusters of target loci alleles of the same direction of fitness effects tend to segregate together, i.e. positive linkage disequilibria arises. This effect increases the range of adaptive frequency oscillations at a target locus, which promotes genetic diversity, provided selection is not too strong.

To make these qualitative conclusions explicit, we present stability analysis of the equilibrium recombination rate between the modifier and target locus, in the infinite-population limit. We then present simulation results on the evolution of the recombination rates, both between the plasticity and the target loci and among comodulated target loci, in the finite populations.

### Stability Analysis

The genomic storage effect leads to the evolution of recombination between the plasticity and the target locus across all of the examined periods of environmental variation (Figure 2) – i.e. in all cases we find evolution to a stable, non-zero rate of recombination, *r*.* The optimal recombination rate *r*\*, that is relatively more fit than all other rates, increases as the period of environmental oscillations (*C*) decreases, similar to what has been observed in other models of fluctuating epistasis (e.g. Charlesworth 1976, Sassaki and Iwasa 1987, Peters and Lively 1999). A non-recombinant modifier is outcompeted by a range of recombination rates when the period is short (*C* = 10), but it can co-exist with modifier alleles for recombination as the environmental period increases. In the absence of an allele encoding the optimal rate *r*\*, two alleles coding for a different recombination rates can coexist in proportions such that on average the population still recombines at rate roughly equal to *r*\*. In other words, in the absence of the optimal recombination rate overdominance may arise at the recombination modifier locus.

**Figure 2.**
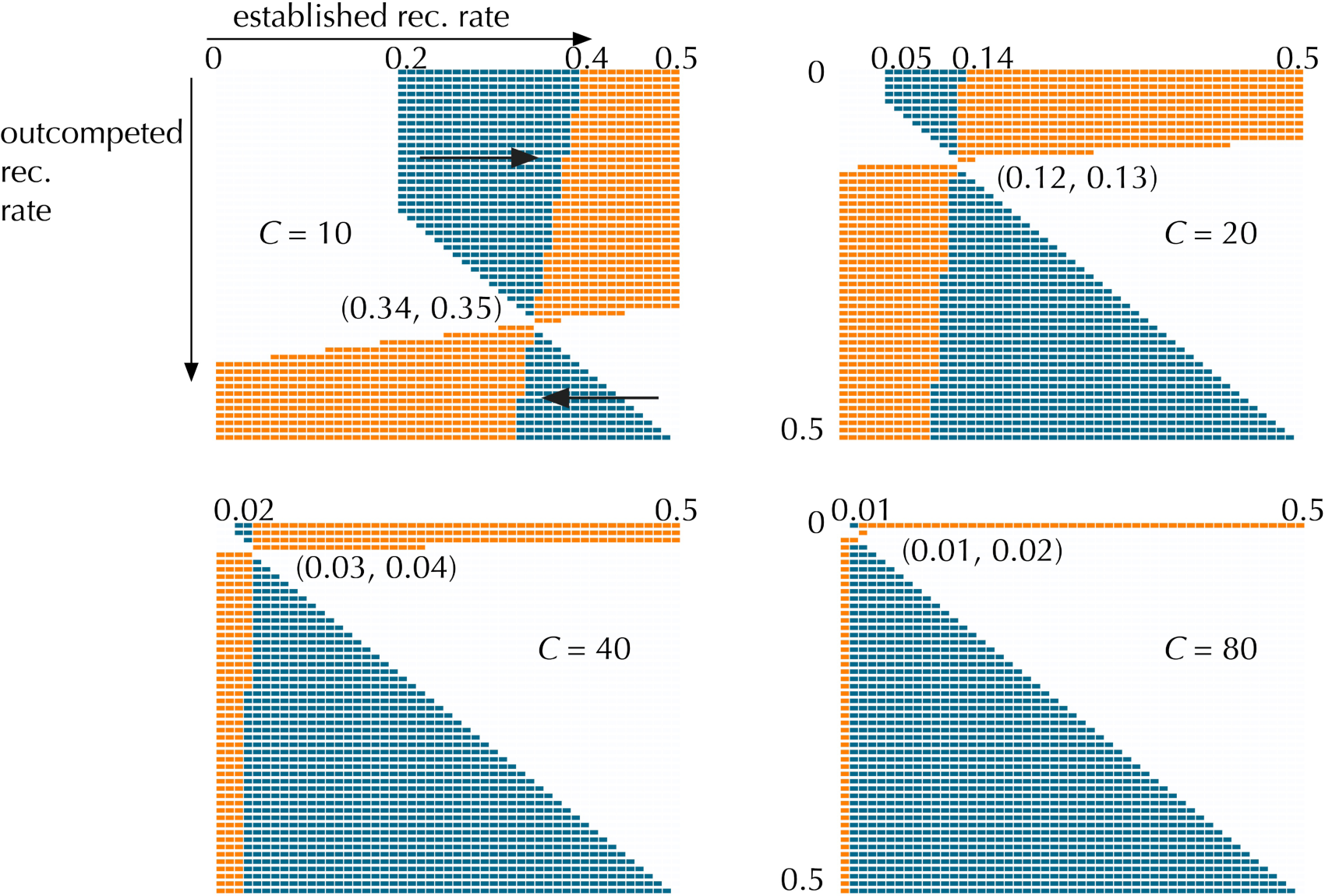
Stability analysis of the recombination rates between the plasticity and the target locus under the genomic storage effect. Each panel indicates a fitter established recombination rate (blue cells), or two balanced rates (orange cells) given a pairwise comparison by a deterministic stability analysis with *C* = 10, 20, 40, or 80. White cells imply that no plasticity-target polymorphism was present at equilibrium. All the blue cells above the diagonal imply evolution towards higher recombination rate, while the blue cells bellow the diagonal mean evolution towards lower recombination rates (see arrows in the first panel). For example, under *C* =10 a non-recombining rate will be outcompeted by all the rates ranging from 0.2 to 0.39, but it will coexist in balanced polymorphism with the rates in the range 0.4 to 0.5. The optimal recombination rate falls within the range given in the brackets. *s*_max_ = 0.1.

Interestingly, stable polymorphic equillibria at both the plasticity and the target locus occur over a wide range of recombination rates, particularly as the environmental period increases. Note that the inferred optimal recombination rates represents a lower bound on the potential optimal rates for each periodicity, because we assumed the free recombination between the recombination modifier and the plasticity-target haplotype (see Charlesworth 1976, Sassaki and Iwasa 1987 or Methods/Stability Analsys).

A natural question is what is the source of selection on a recombination modifier allele in our model? The modifier allele does not itself code for a phenotype, but it is indirectly selected due to the rate at which it produces, and finds itself associated with, selected plasticity-target haplotypes. The relative fitness of a recombination allele (*r*_2_) to the competing recombination allele (*r*_1_), over the period of fitness oscillations (*C*), is given by

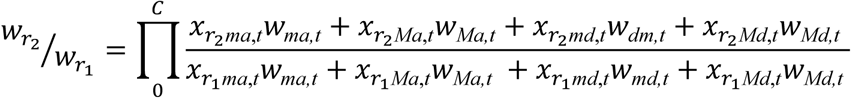

The relative frequencies of the haplotypes in the numerator and denominator of the expression above determine the selective outcome at the recombination locus. For example, consider the selective dynamics over one period of fitness oscillations. Within a season (a sequence of environments of the same direction of selection), selection promotes associations between beneficial allele and a non-plasticity allele, and between the detrimental allele and the plasticity allele (see Gulisija et al. 2016 for details) – a positive linkage disequilibria arises as alleles are paired such to increase their fitness. As the seasons change and the detrimental allele becomes advantageous and vice versa, the newly advantageous allele is associated with the plasticity allele that hinders its increase in frequency while the newly detrimental allele is fully exposed to the effects of negative selection. Thus, the change of seasons results in negative linkage disequilibrium. Recombination results in the increase in frequency of underrepresented fitter haplotypes. An allele for higher recombination rate will increase the proportion of fitter plasticity-target haplotypes much quicker than the one coding for a lower rate of recombination, and hence higher recombination allele will increase its relative fitness. However, this also eventually results in the change of sign in LD, i.e. there will be excess of haplotypes containing allele combinations that maximize fitness, which selection will favor not to decouple. Then, the higher recombination rate will become selected against. It is the balance between these two forces that determine the optimum recombination rate *r*\*. In particular, with longer environmental periods *C*, populations spend more time near equilibrium where a recombination-reducer is favored. It is also evident that a linked recombination modifier allele will gain more selective advantage as it will form a stronger association with the fitter subpopulation – linkage disequilibrium between the recombination modifier and the plasticity locus dissipates at rate = *R* per generation. In support of these stability analyses, we find that in every case where we observe a fixation *r*_2_ we find its fitness relative to that of *r*_1_ (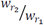) is uniformly greater than one, irrespective of the starting frequencies at the any of the loci, and conversely its relative fitness less than one whenever it perishes. In the cases where two recombination rates coexist, the 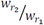 < 1 when the frequency of *r*_2_ allele exceeds the equilibrium frequency, but 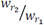 > 1 when it is lower than equilibrium frequency.

### Evolution of recombination in finite populations

In finite populations, genomic storage also leads to the evolution of recombination between a plasticity modifier and its target locus, across a wide range of environmental periodicities (Figure 3). We find that the stationary distribution of the recombination rate coded by an unliked modifier allele subject to recurrent mutation closely agrees with the optimal recombination rate predicted by stability analysis in an infinite population. Also, the peaks of the stationary distribution of the recombination rate (equilibrial rate) at a given periodicity are very close to each even when the strength of selection at the target locus is varied (*s*_max_ *vs. s*_max_/2); thus the evolved recombination rates are robust to the strength of selection in predictably changing environments. With stronger selection, the range of stationary distribution is narrower than under the weaker environmental effect. Although recombination always evolves when environmental periods are short, the stationary distributions for long environmental periods (*C* ≥ 40) include significant probability mass on non-recombinant modifiers (i.e. *r* ~ 0). This too is reflected by the stability analysis in an infinite population, where the non-recombinant type can stably coexist with a large range of positive recombination rates.

**Figure 3.**
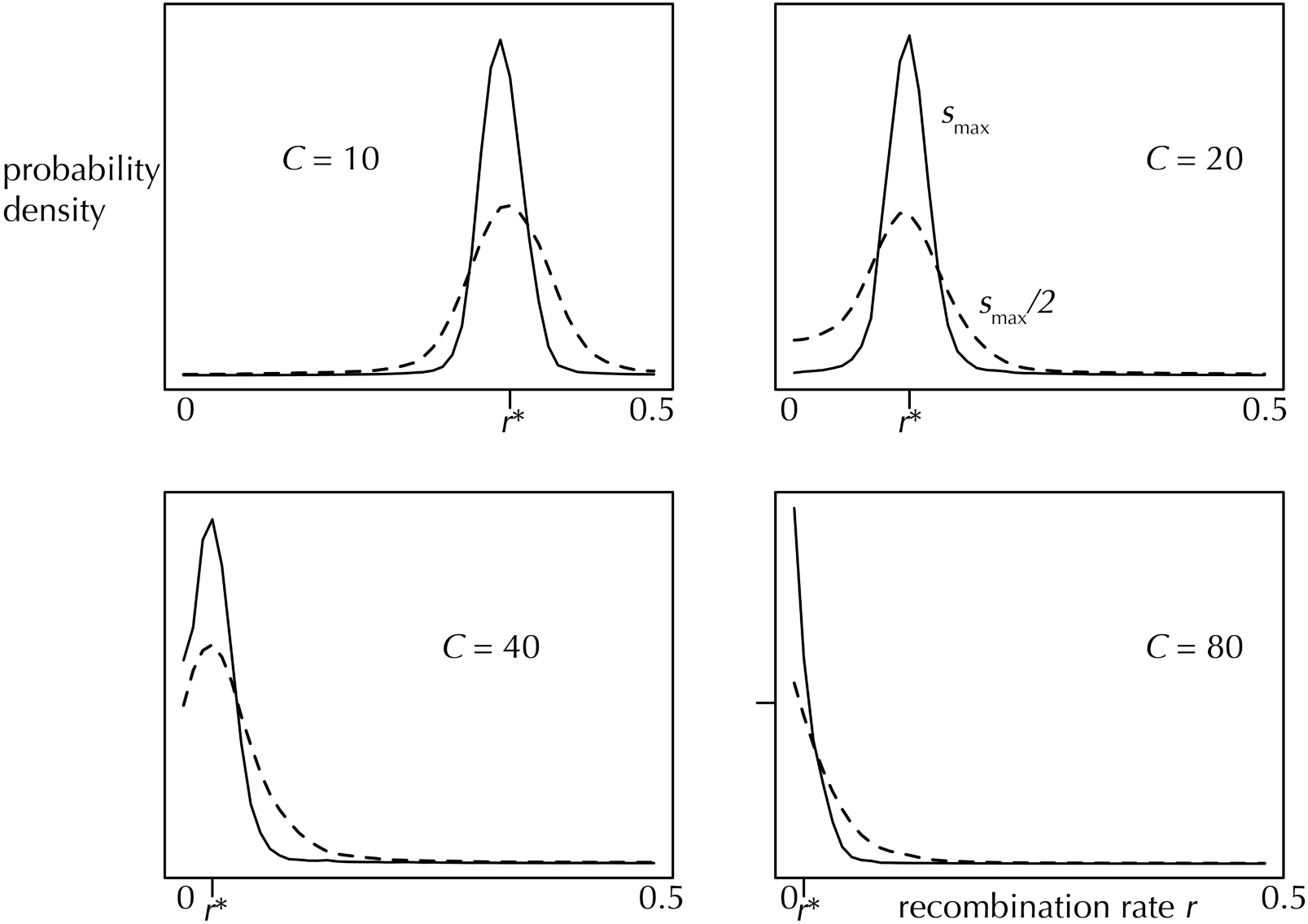
Stationary distribution of the recombination rate *r* between the plasticity and the target locus, encoded by an unlinked recombination modifier subject to recurrent mutation in a population of *N* = 25,000, with *C* = 10 and *s*_max_ = 0.5, *C* = 20 and *s*_max_ = 0.25, *C* = 40 and 80 and *s*_max_ = 0.15 (broken line indicates results with weaker selection, *s*_max_/2). The tick on the horizontal axis denotes the optimal recombination rate (*r*\*) predicted by the deterministic stability analysis. *Nμ* = 0.1. Simulations were run for a 100*N* burn-in generations and the stationary distribution was recorded over the next 100*N* generations. When the strength of selection at the target locus is reduced (by a factor of two, dotted lines), the distribution of recombination rates becomes slightly more broad, but remains centered around the optimal rate.

As before, these results on stationary distributions are conservative lower bounds on the evolution of recombination rate *r*, because all simulations assumed an unlinked recombination modifier (*R* = 0.5). The equilibrial recombination rate is expected to increase if the recombination modifier is linked to the plasticity – target haplotype. Indeed, additional simulations showed larger equilibrial rates when the recombination modifier is flanked by the plasticity and target locus or when it is closely linked to the plasticity locus (*R* = 0.01), than when *R* = 0.5. When the modifier is flanked by the two loci, the equilibrial rate is more than doubled for *C* ≥ 40, since here recombination locus is more closely linked to the selected locus then with *C* ≤ 20 (results not shown). When the recombination modifier is closely linked to the plasticity modifier (*R* = 0.01), free eqilibrial recombination rate (~ 0.5) evolves with *C* = 10, and equilibrial rate is ~ 0.285 for *C* = 20, ~ 0.135 for *C* = 40, and ~ 0.075 for *C* = 80 (Supplementary Figure S2.), showing evolution of recombination in non-recombining population across all periodicities under this scenario.

All the simulations reported above assumed strong selection at the target locus (large *s*_max_) coupled with *N* = 25,000. We expect that the effect extends to the larger populations with lower absolute strength of selection since Gulisija et al (2016) argued that the genomic storage effect depends on the product *Ns*_max_. To verify this, we conducted a two-allele simulation in larger populations, where the optimal recombination rate estimated by infinite-population stability analysis was introduced to a nonrecombinant population. These finite population simulations showed not only that the recombination evolved more readily in larger populations with the same *s*_max_ (Supplementary Figure S4, compare left and middle panel), but more so if the recombination modifier is linked to the plasticity-target sequence. Furthermore, while the effect is weakened by reduction in the plasticity effect, *p*, (Figure 4, right panel), since genomic storage effect decreases with relative plasticity effect (Gulisija et al 2016), recombination still evolves when *p* is small.

**Figure 4.**
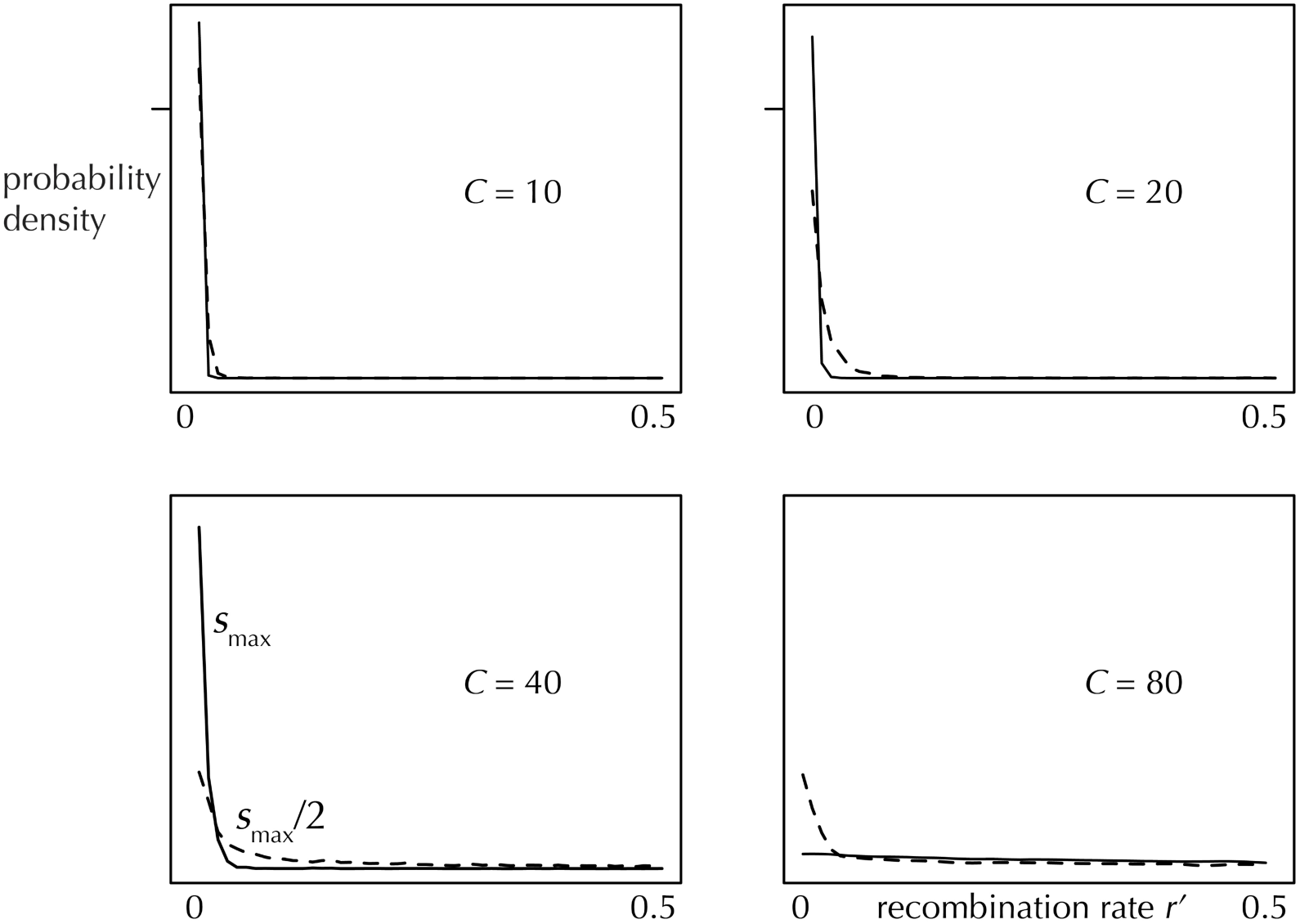
Stationary distribution of the recombination rates between two co-modulated target loci, when the initial target locus starts at the optimal distance to the plasticity modifier loci and a new target locus is introduced downstream away from the plasticity modifier-target haplotype. *N* = 25000 and *C* = 10 with *s*_max_ = 0.5, *C* = 20 with *s*_max_ = 0.25, *C* = 40 and 80 with *s*_max_ = 0.15 for each of the target loci (broken line indicates *s*_max_/2), with *Nμ* = 0.1. Simulations were run for a 100*N* burn-in generations and the stationary distribution was recorded over the next 100*N* generations (to a 2 decimal place precision).

### Evolution of clustering between co-modulated target loci

The genomic storage effect in periodic environment leads to the evolution of reduced recombination, and eventually complete linkage (*r*’ ~ 0), between target loci comodulated by a plasticity modifier. Irrespective of whether the target loci are introduced sequentially or simultaneously, and irrespective of the initial position of the newly introduced target locus or of the location of their recombination modifiers, we find evolution of reduced recombination rates *r*’ among the target loci (i.e. clusters), positive linkage disequilibrium between the loci, and higher levels of diversity (provided the selection is not too strong) and increase magnitude of fitness oscillations at each locus compared to a single-target case. Notably, selection towards reduced recombination among co-regulated target loci seems stronger, resulting in narrow stationary distribution, than selection for the recombination between plasticity and target locus (Figure 4), given the same *Nμ*, *C*, *P* and *s*_max_ for each locus. Interestingly, when the first target locus is controlled by a linked recombination modifier (*R* = 0.01) and the second target locus by an unlinked recombination locus, both loci cluster to the recombination distance with the plasticity locus that is equilibrial under the linked recombination modifier. This occurs because positive LD between the two target loci, which is generated by the genomic storage and reduced recombination, leads to the clustering between two target loci (reduction principle) despite the fact that the two target loci should gravitate to different recombination distances from the plasticity modifier. The clustering of target loci results in a stationary distribution of *r*’ that is almost indistinguishable from that when loci are introduced sequentially, where they gravitate to the same recombination distance from the plasticity modifier (Supplementary Figure S3). The stationary distribution of recombination rates *r*’ is sharply decreasing convex function, more so with stronger selection (*s*_max_ = 0.5, 0.25, or 0.15 for *C* = 10, 20, or 40) where the mean stationary recombination rate is smaller than 0.01, irrespective of environmental periodicity. With weaker selection (*s*_max_ half as large) expected *r*’ is increasing with period to as large as 0.14 (*C* = 80)(Figures 4 and S3). Clustering of target loci was not observed with *C* = 80 and *s*_max_ = 0.15 because in that regime strong selection at clusters of ancestral-ancestral (*a-a*) or derived-derived (*d-d*) alleles pushes alleles to the fixation/loss boundaries and removes polymorphism at the target loci, which is required for the genomic storage effect to operate and thus for the evolution of recombination.

As noted above, genomic storage generates positive linkage disequlibrium between the target loci, which fosters reduced recombination between those sites (recombination decreases because the fittest haplotype is already overrepresented in a population). Hill and Robertson (1966) showed that negative LD could arise even in the absence of epistasis as directional selection quickly fixes/removes combinations of co-adapted/maladapted alleles in finite populations. Under the genomic storage, we find that the balancing selection, as opposite to directional effect, actually maintains strongly selected clusters – that is, haplotypes with largest variance in fitness (ancestral-ancestral or derived-derived), while mismatched haplotypes perish in finite populations. Therefore, we find a notable excess of *a-a* or *d-d* haplotypes than expected in the absence of epistasis between the two loci. With stronger selection and shorter periods, we observe more than three-fold more *a-a* and *d-d* haplotypes than expected in the absence of epistasis.

#### Diversity begets diversity

The genomic storage effect postulates that diversity at the modifier locus promotes diversity at the target locus and vice versa (Gulisija et al 2016). Here, we have shown that this dynamic also hold among co-modulated target loci. That is, a target locus is more likely to be in the intermediate frequency range (minor allele frequency at least 0.1) if it is controlled by the same plasticity locus as another polymorphic target locus, then in the absence of the second polymorphic target locus. This phenomenon occurs because the compound fitness effect increases with the creation of *a-a* or *d-d* haplotypes. Since the strength of the genomic storage effect increases with the strength of selection (provided selection is not so strong as to drive either allele at the target site to fixation within one environmental period), we observe higher levels of diversity at both of the target loci, as compared to the diversity observed at a single polymorphic target locus. Thus, clustered target loci are more likely to remain polymorphic than in the absence of clustering or compared to a single target locus.

#### Evolution of supergenes

Clustering of more than two loci that contribute to the same phenotype does not arise as readily as clustering of just two target loci. Using the same parameters as in the previous section, we find that clustering is unlikely to evolve *de novo* when recombination rates among target loci evolve simultaneously, except for very mild clustering of three target loci under the strongest selection, *s*_max_ = 0.25 (data not shown). However, if the target loci are initiated without recombination, a closely linked sequence of polymorphic alleles acting in unison, i.e. supergenes, will arise despite mutation for recombination at all of the recombination modifier loci and despite mutation at all of the target loci. Therefore, under the genomic storage, supergenes appear unlikely to evolve in the absence of initial proximity to an existing cluster; but they can easily arise the through tandem duplication of target loci.

Indeed, in the sequential model of acquiring new target loci we recover evolution of three-loci clusters and similar stationary distributions of the recombination rates among them as in the two-locus model, when the initial two target loci are initiated very close to each other (initial recombination between first and second target = 0.001, but between second and third ~ U[0,0.5], with *s*_max_ = 0.075). Subsequently the supergenes sequentially grow, including up to *n* = 8 target loci, the maximum number we examined. There, at equilibrium at least 60% of target loci haplotypes occur in the form of all co-segregating *a* alleles or all *d* alleles. As supergenes are created and expanded, the magnitude of frequency oscillations over the period of environmental variation increases.

## DISCUSSION

The Red Queen hypothesis provides a reasonable argument for the evolution of recombination under changing environments, but it is limited to species that co-evolve with an antagonistic species, such as parasites. This study introduces a novel scenario for the evolution of recombination under changing environments that does not require antagonistically interacting species or a constant influx of mutations: the genomic storage effect due to phenotypic plasticity. Gulisija et al (2016) demonstrated the genomic storage effect promotes balanced polymorphism across a range of parameters, including the variation in the strength of benefit or cost of plasticity, the period of environmental change, and even in the presence of random environmental perturbations. However, the polymorphism supported by genomic storage is limited to a relatively large population sizes or strong selection pressures. Nonetheless, large population sizes are not uncommon for many organisms evolving in the periodic environments, such as seasonally evolving organisms (Winkler et al. 2008). Moreover, empirical studies have reported large allele frequency oscillations under temporally varying selection (Lynch 1987; Cain et al. 1990; Turelli et al. 2001), even at many loci simultaneously (Bergland et al. 2014). Thus, phenotypic plasticity may provide a plausible mechanism for the evolution and recombination in periodic environments, and, unlike in the previous studies, even when a single locus codes the phenotype under selection.

The genomic storage effect not only promotes recombination between a plasticity modifier and its target locus, but it simultaneously suppresses recombination among two co–modulated target loci, producing clusters with aligned allelic effects. Previous research into reduction of recombination between polymorphic loci suggests that this phenomenon would be unlikely in the absence of initial physical linkage (Charlesworth and Charlesworth 1975). Surprisingly, under the genomic storage, the clustering of two target loci can readily arise independent of initial physical linkage between the loci and in the absence of epistasis between them. As genomic storage maintains polymorphism, the two loci gravitate to a same recombination distance from the plasticity locus that modulates their effects. Furthermore, genomic storage in finite populations promotes positive linkage disequilibrium between co-modulated loci, which further promotes reduced recombination among target loci. The clustered alleles act in synchrony to magnify the range of adaptive frequency oscillations at each locus, and to increase diversity at the both loci despite the lack of epistatic interactions between them.

While two target loci under the genomic storage will evolve to be clustered independent of initial linkage, the evolution of more than two loci clustering only occurs in a sequential fashion: when existing loci are clustered, and new locus evolves to be tightly linked as well, such that a supergene can emerge sequentially. Here, again, selection generates strong positive linkage disequilibrium within a cluster, which promotes polymorphism. The joint effect of linkage disequilibrium and storage on creation of supergenes suggests that other forms of storage effects might favor supergenes, such as storage due to population subdivision (Gulisija and Kim 2015). Whether or not such scenario is likely in nature and whether other models of storage effect are more likely to give rise to supergenes, however, is subject for future research.

Our study highlights a role for recombination modification in the maintenance of genomic variation in variable environments. Polymorphism under the genomic storage effect persists only in the presence of recombination between the plasticity modifier locus and its target locus. Such recombination will naturally evolve, we have shown, and then subsequently promote balanced polymorphism at both loci. If there are multiple target loci, they will tend to cluster together and exhibit positive LD, compounding their additive effects on fitness, and further enhancing the strength of the genomic storage effect. Thus, the maintenance of diversity by genomic storage is tightly linked to evolution of recombination rates.

Finally, this study points to the importance of phenotypic plasticity in shaping the recombination rates across genome, and the diverse effects it may have on genetic architecture in periodic environments.

## SUPPLEMENTARY FIGURES

**Figure S1.**
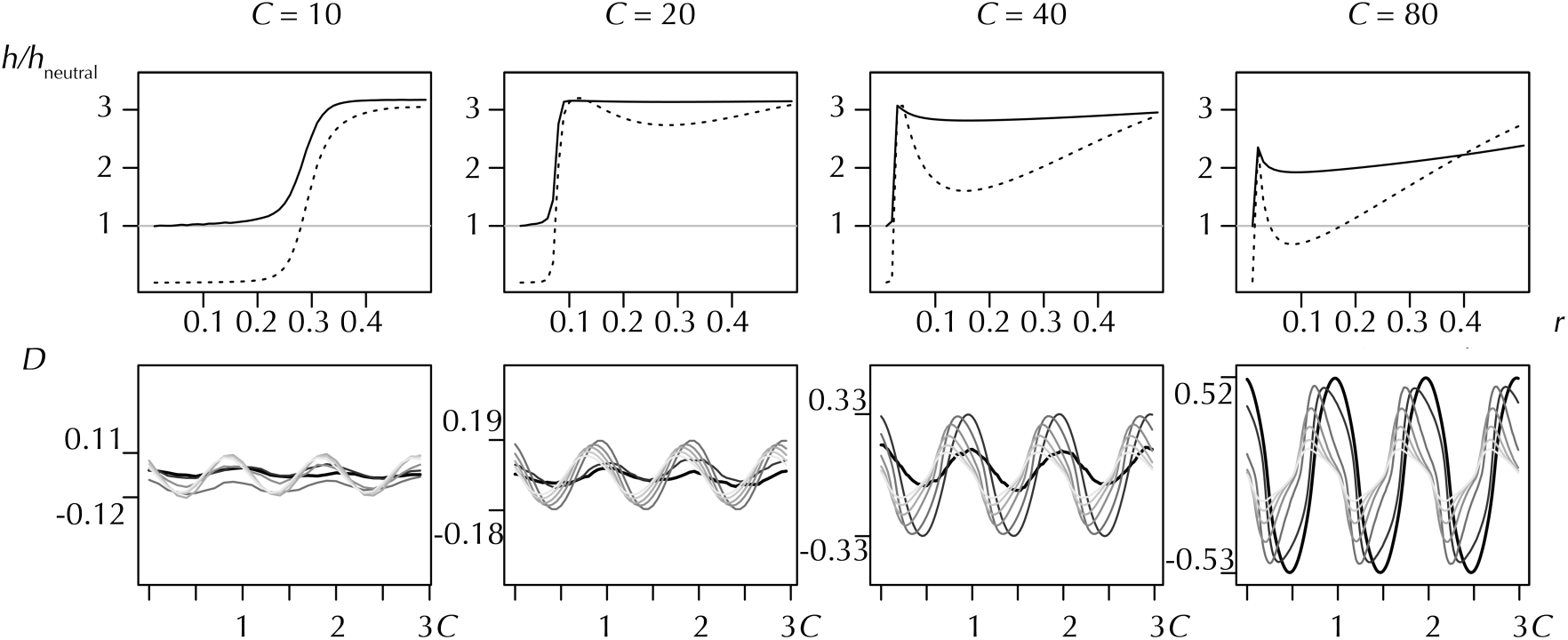
Genomic storage effect generates both balanced polymorphism and cycling linkage disequilibrium. Top row shows the levels of heterozygosity (*h* = 2*p*(1−*p*), where *p* is the frequency of an allele) relative to that expected under neutrality, at the target (solid line) and the plasticity modifier locus (dotted line) across the range of recombination rates [0, 0.5], in a population of size of *N* = 25000 and the selection strength *s*_max_ = 0.1 under the period of selection *C* = 10, 20, 40 and 80 generations (left to right), and the recurrent mutation rate *Nμ* = 0.1 at both of the loci. Bottom row shows normalized linkage disequilibrium patters (*D* = *D*_observed_/*D*_max_, Lewontin 1964) for recombination rates of 0.01, 0.03, 0.1, 0.2, 0.3, 0.4, or 0.5 (ranging from darkest to lightest shade of grey, with 0.01 being ticker line) over the 3 cycles of periodic selection (*C*). Here, recombination rate is assumed intrinsic and does not evolve, for details please see the Model section.

**Figure S2.**
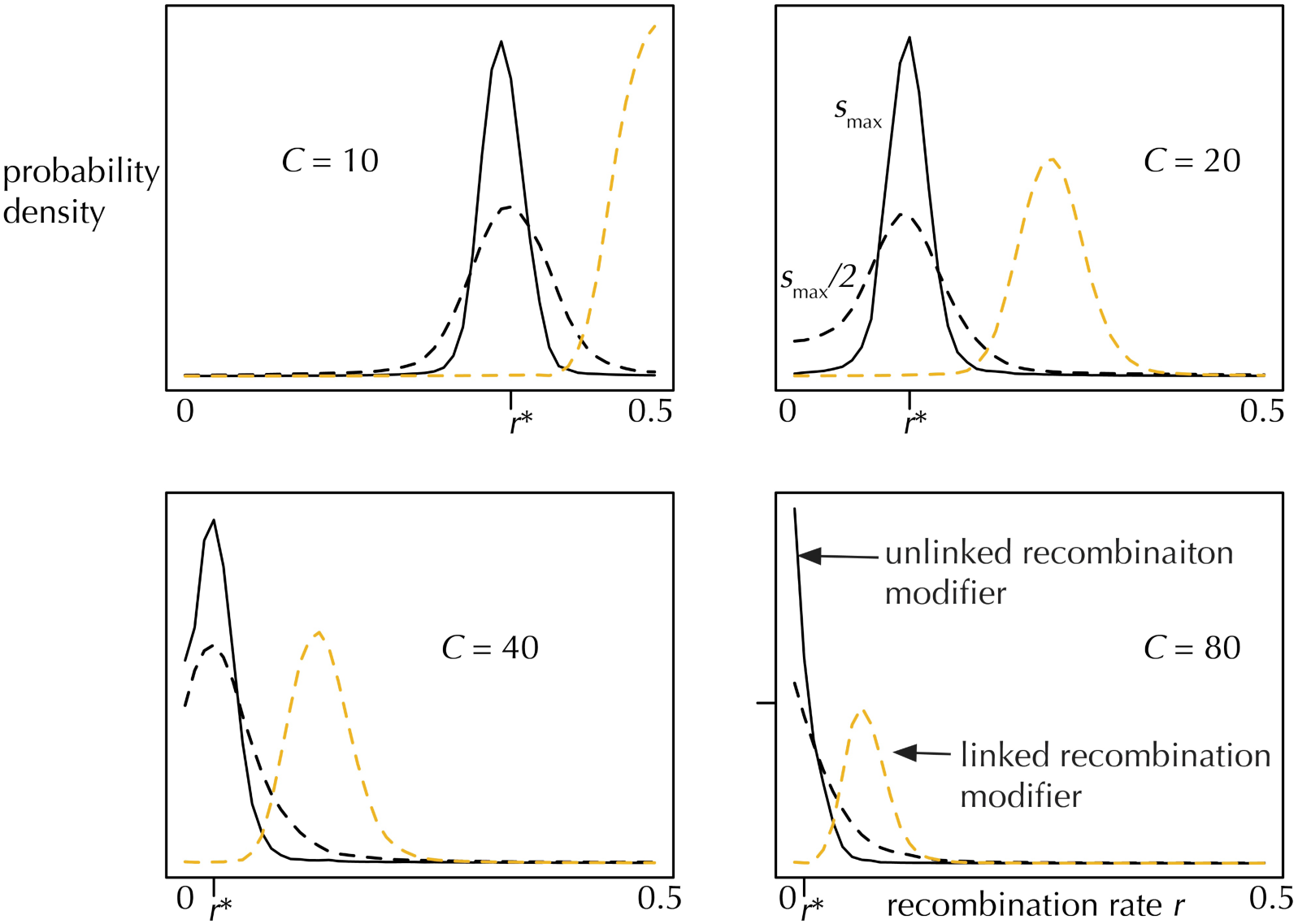
Stationary distribution of the recombination rates between the plasticity and the target locus, in a population of *N* = 25000, with *C* = 10 and *s*_max_ = 0.5, *C* = 20 and *s*_max_ = 0.25, *C* = 40 and 80 and *s*_max_ = 0.15 (broken line indicates *s*_max_/2) assuming the rate is coded by a recombination modifier that is unlinked (black) or linked (with *R* = 0.01, gold) to the plasticity-target sequence. A tick mark on the horizontal line points to the optimal recombination rate (*r*\*) obtained in the deterministic stability analysis with *s*_max_ = 0.1. *Nμ* = 0.1. The distribution is gathered over the 100*N* generations past the 100*N* generations of burn-in period.

**Figure S3.**
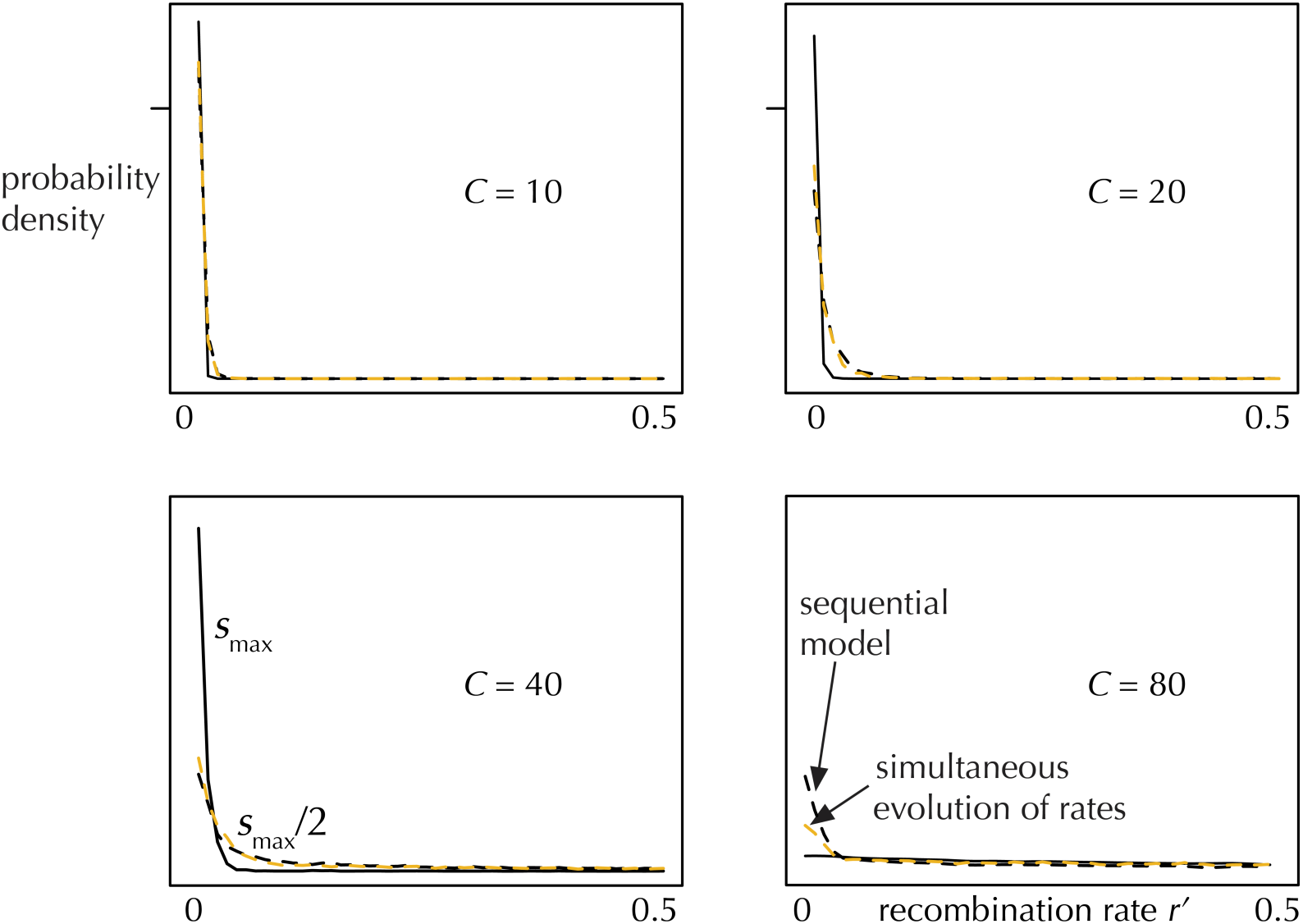
Stationary distribution of the recombination rates between the two co-modified target loci, where the initial target locus stars at the optimal distance to the plasticity modifier (sequential model, in black), or where both recombination rate between the plasticity modifier and the initial target locus evolve (due to linked recombination locus with *R* = 0.01), and between two target loci (due to unlinked recombination locus, *R* = 0.5), given in gold, evolve simultaneously. *N* = 25000 and *C* = 10 with *s*_max_ = 0.25, *C* = 20 with *s*_max_ = 0.125, *C* = 40 and 80 with *s*_max_ = 0.075 for each of the target loci (broken line indicates *s*_max_/2), with *Nμ* = 0.1. The distribution is gathered over the 100*N* generations past the 100*N* generations of burn-in period.

**Figure S4.**
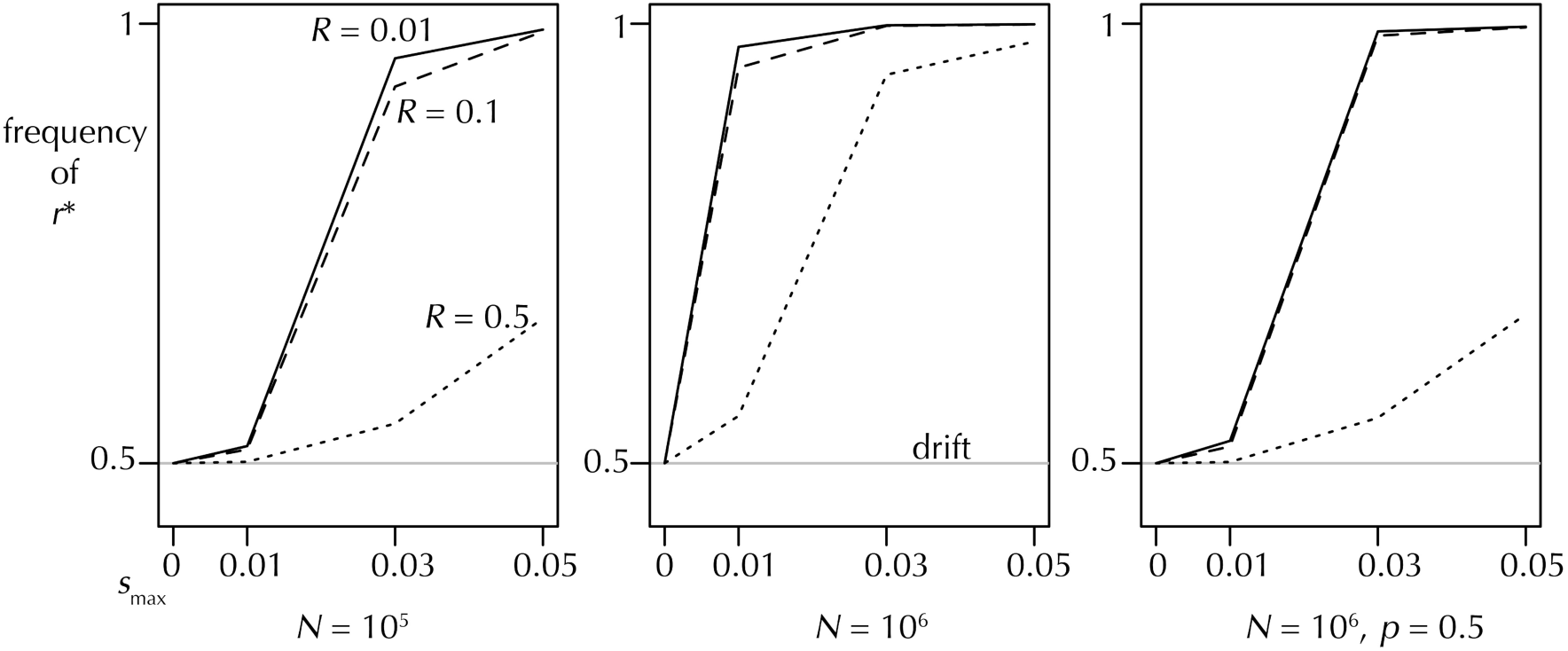
Evolution of recombination rates is more likely under larger population size or when the recombination modifier locus is linked to the plasticity-target sequence. Figure shows equilibrium frequencies of *r*\* in initially non-recombining population under the genomic storage effect. *C* = 20, *Nμ* = 0.1, and *p* = 1 in the first three panels. The frequency is measured at equilibrium, after the 100*N* generations of burn-in, assuming no linkage (*R* = 0.5), weak (*R* = 0.1) and strong linkage (*R* = 0.01) between the recombination locus and the plasticity-target sequence. 40000 simulation runs were conducted.

